# Flow cytometry allows rapid detection of protein aggregates in cell culture and zebrafish models of spinocerebellar ataxia-3

**DOI:** 10.1101/2021.03.09.434364

**Authors:** Katherine J Robinson, Madelaine C. Tym, Alison Hogan, Maxinne Watchon, Kristy C Yuan, Stuart K Plenderleith, Emily K Don, Angela S Laird

**Affiliations:** Centre for Motor Neuron Disease Research, Department of Biomedical Sciences, Faculty of Medicine, Health and Human Sciences, Macquarie University, Sydney Australia

**Keywords:** Spinocerebellar ataxia-3, Machado Joseph disease, hereditary spinocerebellar ataxias, neurodegenerative disease, flow cytometry, proteinopathy, insoluble protein species, protein aggregates

## Abstract

Spinocerebellar ataxia-3 (SCA3, also known as Machado Joseph Disease), is a neurodegenerative disease caused by inheritance of a *ATXN3* gene containing a CAG repeat expansion, resulting in presence of a polyglutamine (polyQ) repeat expansion within the encoded human ataxin-3 protein. SCA3 is characterized by the formation of ataxin-3 protein aggregates within neurons, neurodegeneration, and impaired movement. In this study we have identified protein aggregates in both neuronal-like (SHSY5Y) cells and *in vivo* (transgenic zebrafish) models expressing human ataxin-3 protein containing polyQ expansion. We have adapted a flow cytometric methodology, allowing rapid quantification of detergent insoluble forms of ataxin-3 fused to a green fluorescent protein. Flow cytometric analysis revealed an increased number of detergent-insoluble ataxin-3 particles in cells and zebrafish expressing polyQ expanded ataxin-3 when compared to cells and zebrafish expressing wildtype human ataxin-3. Interestingly, a protein aggregation phenotype could be detected as early as two days of age in transgenic zebrafish, prior to the onset of a detectable movement impairment at 6 days of age, suggesting protein aggregation may be an early disease phenotype in SCA3. Further, treatment of SCA3 cells and transgenic zebrafish with compounds known to modulate the activity of the autophagy protein quality control pathway altered the number of detergent-insoluble ataxin-3 particles detected by flow cytometry. We conclude that flow cytometry is a powerful tool that can be harnessed to rapidly quantify ataxin-3 aggregates, both *in vitro* and *in vivo*, and can be utilised to screen and compare potential protein aggregate targeting therapies.

## Introduction

Spinocerebellar ataxia-3 (SCA3, also known as Machado Joseph Disease or MJD) is a devastating neurodegenerative disease that causes progressive ataxia (loss of balance and co-ordination) and paralysis, as well as impaired speech, swallowing and vision due to the progressive death of neurons in the central nervous system [1]. SCA3 is the most common form of hereditary ataxia found throughout the world (21-28% of autosomal-dominant ataxia) [2-4], affecting approximately 1-5 per 100 000 people worldwide.

SCA3 is caused by inheritance of an abnormal form of a gene called *ATXN3* [5, 6]. The *ATXN3* gene normally contains a repetitious sequence of genetic code (CAG repeat region), which in turn encodes a long string of glutamine (Q) residues (known as a polyQ region) within the ataxin-3 protein. Whilst the normal (wild-type) ataxin-3 protein contains 12-44 glutamine residues within the polyQ region, expression of ataxin-3 with more than 44 glutamine residues produces SCA3 disease in patients [1, 7]. A direct relationship exists between the length of this polyQ region and the severity of SCA3, with patients carrying longer repeat lengths suffering earlier disease onset, more severe disease and earlier death [7-10].

Neuropathological staining of SCA3 patient brain samples often reveals the presence of ataxin-3-positive neuronal intranuclear inclusions [11]. Formation of these ataxin-3 protein aggregates and inclusions is hypothesized to play a role in the neuronal dysfunction and degeneration that occurs in SCA3 [11-13]. Whilst the ataxin-3 protein has been found to function as a de-ubiquinating enzyme [14], ataxin-3 has also been found to sequester other essential proteins into protein aggregates, altering the essential functions of sequestered proteins [15, 16]. Similarly, other proteins containing polyglutamine tracts, such as huntingtin and the androgen receptor, have also been found to aggregate when the polyglutamine tract is expanded, resulting in neurodegenerative diseases such as Huntington’s disease and Kennedy’s disease, respectively [17, 18]. For these reasons, approaches to identify therapeutics that can prevent, or reverse, protein aggregation processes are currently an area of thorough investigation within the field [19].

Whiten et al [20] previously established a flow cytometric approach enabling rapid enumeration of insoluble protein inclusions present in cell culture models, coined FloIT. This approach has previously been used to rapidly quantify the number of detergent insoluble inclusions across three *in vitro* models: cells expressing mutant huntingtin (linked with Huntington’s disease) and in cells expressing the motor neuron disease-linked proteins mutant SOD1 [20] and TDP-43 [21]. Considering that formation of detergent-insoluble protein aggregates is a disease mechanism common to a wide variety of diseases, including Alzheimer’s disease, Parkinson’s disease, polyglutamine diseases such as Huntington’s disease and spinocerebellar ataxias, and even type 2 diabetes [22-26], it is important to determine whether FloIT is also applicable to other proteinopathy disease models. This is especially important to determine when considering proteinopathies within differing cellular compartments. For example, protein aggregates found in models of motor neuron disease and Huntington’s disease are more commonly present within the cell cytoplasm [27, 28], whilst in other diseases, such as spinocerebellar ataxia-3, protein aggregates are more commonly found within the cell nucleus [11, 29].

In the present study, we demonstrate that FloIT can be adapted to quantify the number and size of detergent-insoluble EGFP-fused ataxin-3 particles in neuronal-like (SHSY5Y) cells transiently expressing ataxin-3 containing a short polyQ region length (28 glutamine residues) or long polyQ region length (84 glutamine residues). We also report adaption of this methodology for use with our previously reported transgenic zebrafish model of SCA3 [30]. This adaption is important because zebrafish are an excellent model for moderate to high throughput assessment of drug efficacy, as overexpression of human genes can produce disease phenotypes akin to human disease phenotypes, including impaired movement, within days. Further, potential therapeutic compounds can be diluted into the water the fish live in and easily absorbed [31]. This is the first report of use of such an approach with an *in vivo* model and provides a novel tool to assess efficacy in drug testing studies investigating treatments for proteinopathy diseases.

## Materials and Methods

### SCA3 Cell Culture Models

Human neuroblastoma (neuronal-like) SHSY5Y cells were grown under sterile conditions and maintained at 37°C in a sterile incubator. Cells were grown in DMEM/Nutrient Mixture F12 Ham and supplemented with 10% fetal bovine serum and 5% CO_2_. Cells were seeded into 6-well plates and allowed to grow for 1-3 days until 80-90% confluent. Once confluent, cells were transiently transfected with vectors to drive expression of EGFP fused to human ataxin-3 with a short polyQ repeat length (28Q) or human ataxin-3 with an expanded polyQ repeat length (84Q) or an EGFP only vector control. Cells were transfected with 1 μg of DNA and 1 μL of Lipofectamine 2000 (Thermo Fisher) per well and incubated for 24 hours.

### Confocal microscopy of aggregates in a SHSY5Y cell model of SCA3

For confocal imaging experiments, SHSY5Y cells were seeded into a 24 well-plate and grown on coverslips until reaching 60-70% confluency. Cells were then transfected with 1 μL of Lipofectamine 2000 and 1 μg of DNA per well. At 24 hours post-transfection, culture media was aspirated, and cells were fixed with 4% paraformaldehyde and stained with 4’,6-diamidino-2-phenylindole (DAPI, final concentration 14 μM) to enable identification of nuclei, then mounted using fluorescent mounting media (Dako Omnis, Aglient catalogue # GM304). Confocal images were obtained using a Zeiss LSM-880 confocal microscope (Plan-Apochromat 40x/1.5 oil DIC M27 objective) running Zen Black software (Zeiss, Gottingen, Germany). In order to visualise EGFP expression, an argon laser was used. To visualise DAPI-stained nuclei, a 405 nm laser was used. A total of six z-stack images (13 slices spanning 6 μm) of different fields of view were acquired per coverslip and final images were obtained as maximum intensity projections using ImageJ software or Fiji software. EGFP-positive aggregates were manually counted by a researcher blind to experimental group. The number of aggregates per transfected cell was then calculated by averaging the results from six fields of view per coverslip.

### Preparation of SCA3 cell cultures for FloIT

For flow cytometry experiments, SHSY5Y cells were seeded into 6-well plates and prepared for flow cytometry 24-hours post-transfection. Cultured SHSY5Y cells were imaged with 20x objective using an EVOS microscope (Invitrogen, catalogue number AMF4300) under brightfield and 488 nm lasers to determine transfection efficiency prior to harvesting. Cells were briefly washed in PBS and harvested using 0.5% trypsin-EDTA and pelleted. Pelleted cells were re-suspended in lysis buffer (phosphate buffered saline containing 0.5% Triton-X 100 and complete protease inhibitors [Roche]) and DAPI was added (final concentration 5μM) to allowed enumeration of the number of nuclei. Samples were incubated on ice and protected from light until analysis. All samples underwent flow cytometric analysis within 30 minutes of lysis with Triton-X 100.

### Transgenic SCA3 Zebrafish

The present study utilised a previously described zebrafish model of SCA3 [30]. Within these studies we used embryos resulting from crossing zebrafish driver line mq15:

Tg(elavl3:Gal4-VP16; mCherry) with either line mq16:

Tg(UAS:Hsa.ATXN3_23xCAG-EGFP,DsRed) or mq17:

Tg(UAS:Hsa.ATXN3_84xCAG-EGFP,DsRed), resulting in transgenic zebrafish neuronally expressing both dsRED and EGFP-fused ATXN3 with either short (23Q) or long (84Q) polyQ lengths, respectively. All animal experiments were performed in accordance with the Code and approved by the Animal Ethics Committee of Macquarie University (2016/004) and the Biosafety Committee of Macquarie University (Notifiable Low Risk Dealing: 5974). Zebrafish were housed in a standard recirculating aquarium system maintained at 28.5°C. Zebrafish embryos were screened for fluorescence (EGFP and dsRED) at 1-day post-fertilisation (1 dpf) indicating expression of transgenic lines. Embryos were dechorionated and housed in 6-well plates (15-25 larvae per well, housed at equal densities per group).

### Imaging of aggregates in a transgenic zebrafish model of SCA3

At 6 days post-fertilisation (6 dpf), transgenic zebrafish were anaesthetised with 0.01% Tricaine (Sigma, catalogue number E10521) and embedded in 1% low-melting agarose (Sigma, catalogue number A4018). Confocal imaging was performed using an upright Leica SP5 confocal microscope (40x water submersible objective). An argon laser (39% laser was used to excite EGFP. Z-stacks spanning the depth of the spinal cord (∼10 μm) were acquired and final images were obtained as maximum intensity projections using ImageJ software or Fiji software. EGFP positive aggregates were manually counted by a researcher blind to experimental group.

### Preparation of Dissociated of Zebrafish Larvae for Flow Cytometry

Whole zebrafish larvae (2 or 6 dpf) were euthanised and larvae bodies were manually dissected in E3 media using a scalpel blade. Larvae were enzymatically digested using 0.5% Trypsin-EDTA (500 μL for 2dpf larvae, 1000 μL for 6dpf larvae) for 30 minutes at 37°C. Samples were vortexed frequently to aid digestion. Trypsinisation was stopped by the addition of 1 mL of DMEM cell culture media containing 10% fetal bovine serum. Samples were pelleted and the DMEM/trypsin solution was removed. Samples were resuspended in 200 – 500 μL of PBS containing 0.5% Triton-X and 1 x Red Dot far red solution (Gene Target Solutions, catalogue number 40060) to identify nuclei.

### Drug treatments in SCA3 cell cultures and transgenic SCA3 zebrafish

As a proof of principle, we examined the effect of prior treatment with compounds known to alter activity of the autophagy protein quality control pathway on the number of detergent-insoluble ataxin-3 particles detected using flow cytometry, in SCA3 cell cultures and transgenic zebrafish. For autophagy inhibition experiments, cultured cells were treated with 3-methyladenine (3MA, 5 mM, Cayman Chemicals [catalogue number 13121]) and zebrafish larvae were treated with chloroquine diphosphate (3 mM, Sigma-Aldrich [catalogue number C6628]), two commonly utilised inhibitors of autophagy [32]. To examine the effect of autophagy induction on detergent-insoluble ataxin-3, calpeptin (Caymen Chemicals [catalogue number 14593]), a compound known to induce autophagy in SCA3 zebrafish [30] and suppress protein aggregation in cell culture [33, 34], was administered at doses ranging from 1 -5 μM in cell culture and 25 μM – 100 μM for zebrafish. For drug treatments of SHSY5Y cells transiently expressing human ataxin-3, 3MA or calpeptin treatments were dissolved in culture media (DMEM/Nutrient Mixture F12 Ham and supplemented with 10% fetal bovine serum) and applied immediately prior to the addition of transfection reagents. For drug treatments of transgenic zebrafish, chloroquine diphosphate or calpeptin were diluted in E3 raising media and larvae were placed in drug treatments for a minimum of 24 hours. All drug treatments were dissolved in dimethyl sulfoxide (DMSO) to create stock solutions. Stock solutions were aliquoted and stored at -20°C. Treatment with DMSO alone acted as the vehicle control.

### Flow Cytometry Analysis Approach

Flow cytometry was performed using a Becton Dickson Biosciences LSR Fortessa analytical flow cytometer running FACS DIVA software and maintained according to manufacturer’s instructions (Becton Dickson). A minimum of 20,000 events were captured for experiments involving culture cells and 50,000 events were captured for experiments involving dissociated cells from whole zebrafish larvae. The fluorescence of EGFP-expressing cells was compared to an un-transfected or non-transgenic control sample. Nuclei were identified and quantified based on intensity of UV/infrared fluorescence and particle size (forward scatter). The number of detergent-insoluble EGFP particles, indicating insoluble EGFP-fused ataxin-3 particles, were analysed based on GFP fluorescent intensity and forward scatter [20]. For genotype comparisons, the number of detergent-insoluble EGFP-positive particles was determined using the FloIT equation published by Whiten and colleagues [20]: 100 x (# of Insoluble GFP Particles/ # nuclei x transfection efficiency). This analysis approach allows for comparison of detergent-insoluble particles across different transgenic expression models. For comparison of treatment effects on the same transgenic model, the number of detergent-insoluble EGFP-positive particles per number of nuclei present was calculated and presented as a fold-change relative to the vehicle treated control group. Non-fluorescent microspheres (Thermo Fisher, catalogue number F13838) of a known diameter (2 – 15 μm) were used to equate forward scatter measurements of detected to a precise micron diameter.

### Data analysis

Data analysis was performed using GraphPad Prism (Version 8) software. Group comparisons were made using one-way ANOVA tests, followed by a Tukey post-hoc to identify differences. In cases where only two groups were compared, comparisons were made using a student t-test. All graphs display group mean data ± standard error mean.

## Results

### Expression of human ataxin-3 with expanded polyglutamine repeat in neuronal-like SHSY5Y cells results in the formation of EGFP-positive aggregates

Microscopic analysis of SHSY5Y cells expressing EGFP fused-human ataxin-3 containing 28Q (EGFP-ataxin-3 28Q), EGFP-ataxin-3 84Q or EGFP alone, revealed the presence of more frequent EGFP-positive aggregates within the cells expressing human ataxin-3 containing 84Q than 28Q or EGFP-only vector control (Figure 1A). Manual counting of the number of EGFP-positive protein aggregates and one-way analysis of variance revealed a significant difference in the number of EGFP-positive aggregates across genotypes (*p* < 0.0001). Post-hoc comparisons revealed that more aggregates were present in the SHSY5Y cells expressing EGFP-ataxin-3 84Q when compared to cells expressing EGFP alone (*p* < 0.0001) or EGFP-ataxin-3 28Q (*p* = 0.0002; Figure 1B).

**Figure 1.**
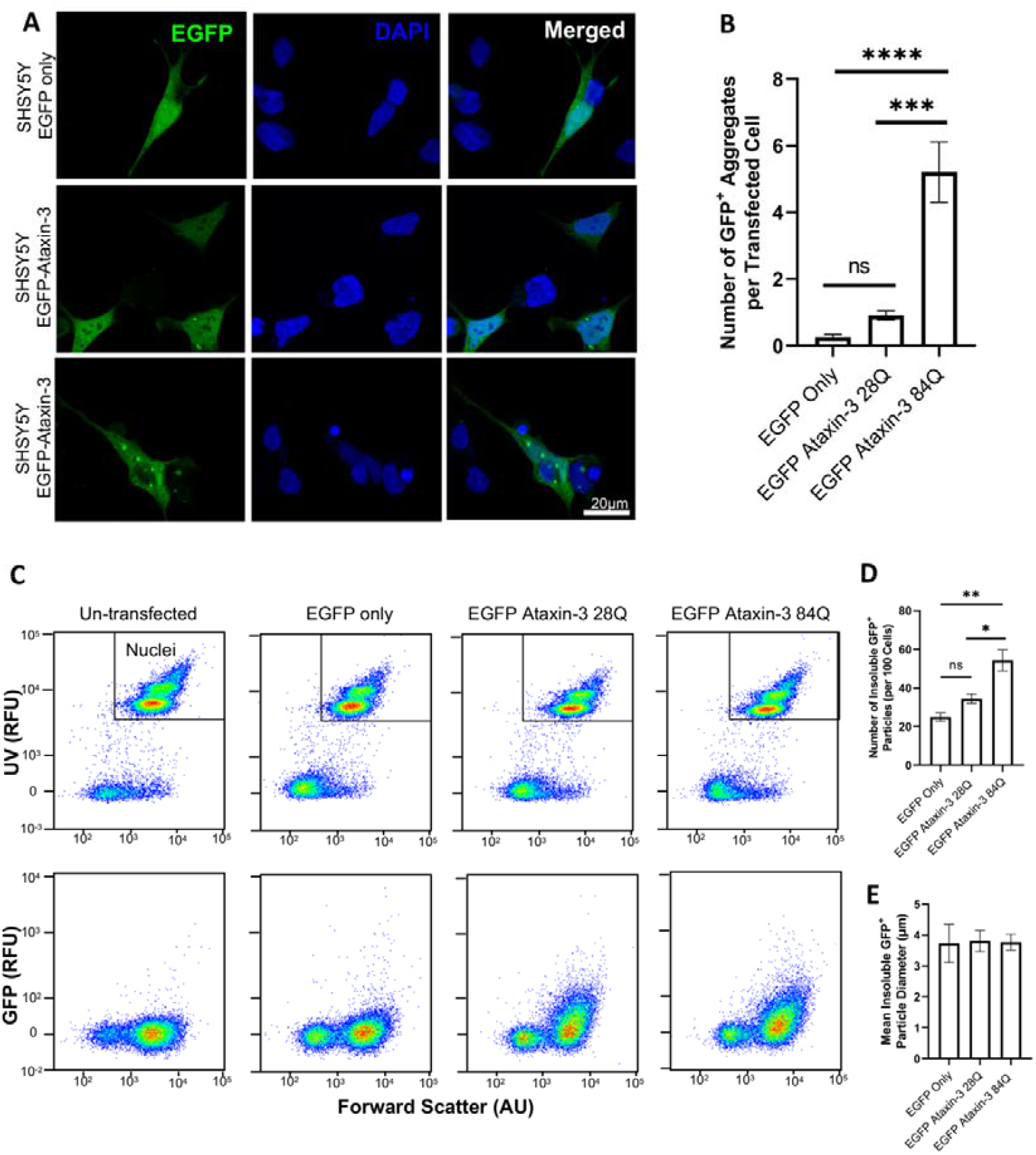
Confocal microscopy and flow cytometry can both be used to detect protein aggregates in SHSY5Y cells transiently expressing EGFP-tagged human ataxin-3. (A) SHSY5Y cells transiently expressing EGFP-fused human ataxin-3 or EGFP alone develop protein aggregates. (B) The mean number of aggregates per transfected cell was calculated by averaging six experimental replicates (different fields of view) per coverslip. Aggregates were more commonly observed in cells transfected with the 84Q repeat expansion than cells transfected with ataxin-3 28Q. (C) Flow cytometry results demonstrating that this method can be used to quantify nuclei and detergent-insoluble EGFP-fused ataxin-3 particles from cell lysates. (D) Cells expressing EGFP-fused mutant ataxin-3 (84Q polyglutamines) develop significantly more insoluble particles when compared to EGFP alone or EGFP-fused ataxin-3 with a short polyglutamine stretch (28 polyglutamines). (E) Detergent-insoluble particle size did not significantly differ across examined genotypes. * indicates p < 0.05, ** indicates p < 0.01, *** indicates p < 0.001 and **** indicates p < 0.0001. Graphs depict mean ± standard error mean, n = 4-9 experimental replicates.

We next validated the rapid quantification of the number Triton-X 100 insoluble aggregates using the previously reported FloIT methodolgy [20]. Transfected cells underwent flow cytometric analysis to quantify the number and size of Triton-X insoluble EGFP-positive particles. Firstly, fluorescent microscopy was used to confirm the number of transfected cells per sample (prior to harvesting). The calculated transfection efficiency of each sample was utilised to calculate the total number of detergent-insoluble particles per sample. Next, the total number of nuclei present within the lysed sample was determined by a DAPI stain for nuclei. The number of DAPI-positive particles was found to be similar in all lysed samples and the total number of DAPI nuclei present in each sample was quantified using population gating based on intensity of UV fluorescence and relative size (forward scatter, Figure 1C). The number of Triton-X 100 insoluble EGFP-positive particles was then identified via gating of particle populations present within cells expressing EGFP constructs but absent in the untransfected control sample [20]. The number of Triton-X insoluble EGFP-positive particles was calculated using the FloIT equation: 100 x (# of Insoluble GFP Particles/ # DAPI nuclei x transfection efficiency) as previously reported by Whiten, San Gil [20]. This equation revealed a statistically significant difference across the analysed genotypes (one-way ANOVA: *p* = 0.0013, n = 4-9 experimental replicates per group), with a greater number of detergent-insoluble particles per 100 cells found in EGFP-ataxin-3 84Q-expressing SH-SY5Y cells when compared to cells expressing EGFP alone (*p* = 0.0022) and EGFP-ataxin-3-28Q (*p* = 0.0118) (Figure 1D). No significant differences were found between the number of detergent-insoluble particles expressed in EGFP alone or EGFP-ataxin-3-28Q (*p =* 0.4328). Additionally, use of non-fluorescent microspheres of known size allowed us to identify that EGFP-ataxin-3-84Q cells harbored detergent-insoluble particles that were similar in size to cells expressing EGFP-ataxin-3-28Q and EGFP alone (Figure 1E).

### Treatment of SCA3 cell cultures with compounds known to modulate autophagic activity altered the number of EGFP-fused ataxin-3 particles detected by FloIT

In order to validate the use of FloIT as a tool to screen the effect of compounds on detergent-insoluble protein species proteinopathy, we treated cells transiently transfected with EGFP-ataxin-3 84Q for 24 hours with 3MA, a known inhibitor of autophagy, and calpeptin, a compound known to induce autophagy and reduce proteinopathy in polyQ disease models. Indeed, we found that 24-hour treatment with modulators of the autophagy protein quality control pathway altered the number of EGFP particles detected by flow cytometry (Figure 2A). We found that treatment with 3MA produced a 1.3-fold increase in the number detergent-insoluble EGFP particles present within cells expressing EGFP-ataxin-3 84Q when compared to vehicle treatment (*p* = 0.002, Figure 2B). Further, analysis of cells transfected with EGFP-ataxin-3 84Q treated with increasing doses of calpeptin, a known autophagy inducer, altered the number of detergent-insoluble ataxin-3 particles detected (*p* = 0.028). Post-hoc comparisons revealed that treatment with 1 μM calpeptin did not significantly alter the number of insolube particles detected when compared to vehicle treatment (*p* = 0.331). In contrast, treatment with 2.5 μM or 5 μM calpeptin produced a statistically significant decrease in the number of detergent-insoluble particles, compared to vehicle treatment (*p* = 0.021 and *p* = 0.031, respectively, Figure 2C).

**Figure 2.**
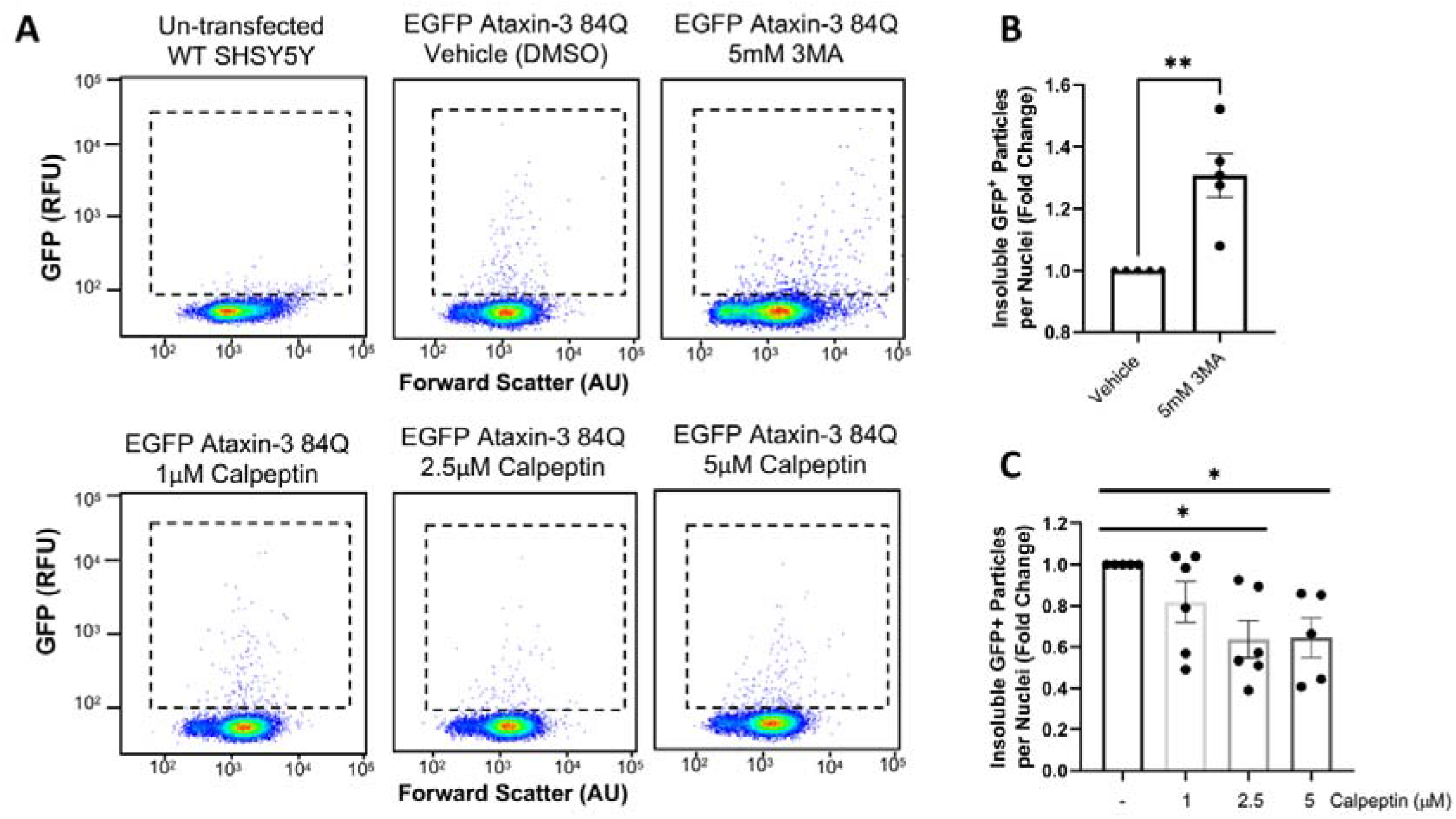
Treatment with autophagy pathway modulators altered the number of EGFP-fused particles present in cells expressing human ataxin-3 with 84 glutamines. (A) Scatterplots reveal a scarcity of EGFP-fused particles in un-transfected control cells and altered numbers of particles following treatment with autophagy modulators (3MA and calpeptin). (B) Comparison of cells treated with vehicle or 3MA (5 mM) revealed a statistically significant increase in detergent-insoluble particles following 3MA treatment. (C) Calpeptin treatment produced reductions in detergent-insoluble particles as detected by FloIT (C). * indicates p < 0.05, ** indicates p < 0.01. Graphs depict mean ± standard error mean, n = 2-6 experimental replicates.

### Expression of expanded human ATXN3 in transgenic zebrafish results in the formation of ataxin-3-positive aggregates

We have previously described a transgenic zebrafish model of SCA3 that overexpresses EGFP-fused human ataxin-3 with expanded polyQ (84Q) tract, that develops impaired swimming behaviour and shortened lifespan [30]. Furthermore, histological analysis of the medulla from adult zebrafish (12 months of age) expressing human mutant ataxin-3 (84Q) evidenced a neuritic beading pattern, that is the presence of ataxin-3 protein aggregates in neuronal neurites,, which was positive for both ataxin-3 and ubiquitin [30]. In this current study we performed confocal microscopy on transgenic zebrafish larvae expressing human ataxin-3 at 6 dpf to examine the rostral spinal cord (Figure 3A). We observed similar levels of overall human ataxin-3 expression in zebrafish expressing ataxin-3 with 23 glutamine residues and ataxin-3 with 84 glutamine residues and observed EGFP-positive protein aggregates in neurons in both transgenic models. Manual counting of the EGFP-positive protein aggregates identified a statistically signficntly higher number of aggregates in larvae expressing EGFP-ataxin-3-84Q than in EGFP-ataxin-3-23Q (*p* = 0.0007; n = 5 zebrafish larvae imaged per ataxin-3 construct, Figure 3B).

**Figure 3.**
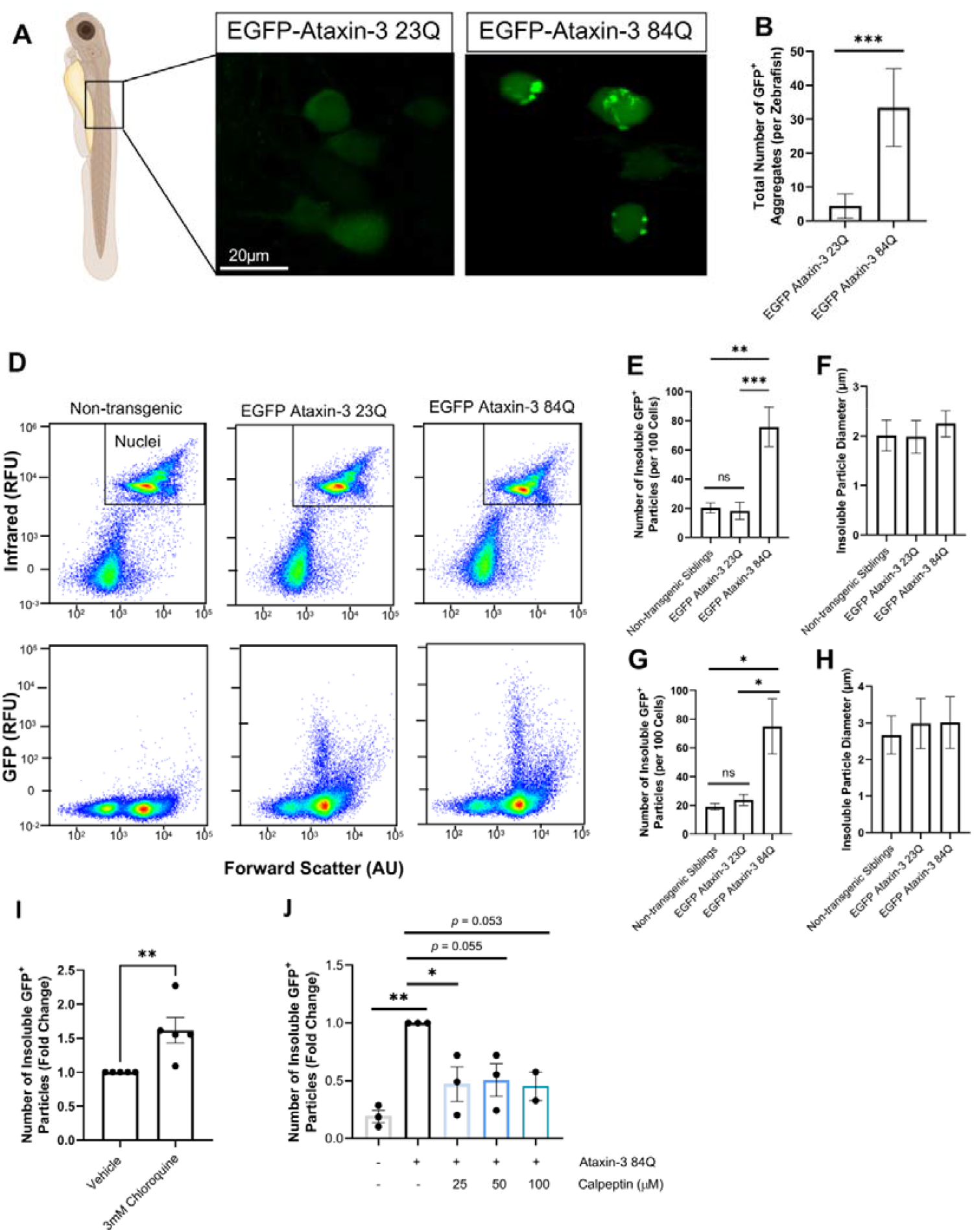
Transgenic zebrafish expressing human ataxin-3 with 84 polyglutamines develop ataxin-3 positive protein aggregates that can be rapidly quantified using FloIT. (A) Confocal microscopy was performed on 6-day old zebrafish larvae expressing EGFP-tagged human ataxin-3 containing a short polyglutamine stretch (28Q) or long polyglutamine stretch (84Q). (B) Manual counting of EGFP aggregates revealed that significantly more aggregates were present in 6 dpf zebrafish expressing EGFP-ataxin-3 84Q then EGFP-ataxin-3 23Q. (C) Scatterplots of flow cytometric populations from dissociated 6-day old zebrafish revealed changes in the number of detected detergent-insoluble particles despite similar numbers of nuclei. (D-E) Zebrafish expressing human mutant ataxin-3 display significantly more detergent-insoluble ataxin-3 particles at 2 days of age; and (F-G) 6 days of age when compared to non-transgenic siblings and zebrafish larvae expressing human wildtype ataxin-3. (H) Treatment with the autophagy inhibitor chloroquine (3 mM) increased the number of detergent-insoluble particles in 2-day post-fertilisation zebrafish expressing mutant ataxin-3; and (I) treatment with the autophagy inducer calpeptin decreased the number of detergent-insoluble particles detected. * indicates p < 0.05, ** indicates p < 0.01, *** indicates p < 0.001. Graphs depict mean ± standard error mean, n = 2-9 experimental replicates.

### Flow cytometry can be utilised to quantify number of ataxin-3 protein aggregates in transgenic zebrafish

We then modified the FloIT approach to allow quantification of the number of detergent-insoluble EGFP-positive protein aggregates within cells dissociated from whole zebrafish larvae (2 or 6 dpf) expressing human EGFP-fused ataxin-3. Following euthanasia, manual dissection and trypsinisation of whole zebrafish larvae bodies resulted in a suspension of dissociated cells that could then be lysed with 0.5% Triton-X for FloIT. Visualisation of lysed particles using an infrared laser (670 nM) revealed similar numbers of red dot labelled nuclei present within samples confirming equal number of cells per sample (Figure 3C). In contrast, visualisation of particles using blue-green laser (525 nM) revealed the presence of detergent-insoluble EGFP-positive populations in the cells dissociated from the zebrafish larvae. These EGFP-positive aggregates were detectable as early as two days post-fertilisation, at which timepoint, one-way analysis of variance revealed a significant difference in the number of detergent-insoluble EGFP-positive particles per 100 cells across examined genotype groups (*p* = 0.0004), with significantly more detergent-insoluble ataxin-3 positive particles in dissociated zebrafish expressing EGFP ataxin-3-84Q compared to non-transgenic siblings (*p* = 0.0012) and EGFP ataxin-3 23Q larvae (*p* = 0.0009, Figure 3E). Interestingly, use of calibrated microspheres revealed that the size of the detergent-insoluble particles was similar across the genotypes and consistent with aggregate diameters found in other *in vivo* models of SCA3 [35] and SCA3 patient brain tissue [29] (Figure 3F).

We also examined aggregate number and size at 6 dpf, an age at which the 84Q transgenic zebrafish present with a swimming phenotype characterised by reduced distance swum [30]. We found that the number of detergent-insoluble ataxin-3 aggregates was consistent at both timepoints examined, with a significant difference across genotype groups still evident at 6 dpf (*p* = 0.0134). Post-hoc analysis revealed that significantly more detergent-insoluble EGFP-positive particles were present in dissociated zebrafish expressing EGFP ataxin-3-84Q compared to non-transgenic siblings (*p* = 0.0214) and EGFP ataxin-3-23Q-expressing transgenic fish (*p* = 0.0343, Figure 3G). The size of detergent-insoluble EGFP-positive particles was again found to be similar across analysed genotypes, with mean detergent-insoluble particle diameter around 3 microns in diameter by 6 dpf (Figure 3H).

As proof of the utility of the FloIT approach to assess efficacy of drug treatments aimed at modifying protein aggregate formation and detergent-insoluble protein species, EGFP-ataxin-3 84Q larvae were treated with 3 mM chloroquine, an autophagy inhibitor for 24 hours. A 1.5-fold increase in detergent-insoluble EGFP particles was observed in chloroquine treated larvae compared to vehicle control larvae from the same clutch (*p* = 0.012, Figure 3I). FloIT also detected a difference in the number of detergent-insoluble EGFP positive particles present following 24-hour treatment with the autophagy inducer calpeptin (*p* = 0.006, Figure 3J). Post-hoc comparisons revealed that treatment with 25 μM calpeptin produced a statistically significant decrease in detergent-insoluble particles when compared to vehicle treatment (*p* = 0.038). Treatment with 50 and 100 μM doses appeared to produced a decrease in detergent-insoluble particles, however these comparisons were not statistically significant (*p* = 0.055 and *p* = 0.053, respectively).

## Discussion

### Flow cytometry approach detects protein aggregates in a cell culture model of SCA3

Here we report that neuronal-like (neuroblastoma) SH-SY5Y cells expressing EGFP-ataxin-3 containing a polyQ expansion (84 polyQ repeats) develop EGFP-positive protein aggregates. These findings align with existing experimental evidence that suggests polyglutamine expanded ataxin-3 can form protein aggregates within cultured murine or human neuroblastoma cells [11, 35, 36] and reports of intraneuronal ataxin-3 positive inclusions in SCA3 brain tissue [11, 13, 29, 37].

Next, we aimed to determine if we could employ the high-throughput flow cytometric analysis approach, FloIT [20], to quantify the number and size of detergent-insoluble ataxin-3 particles in SCA3 cells. This method has been previously utilized to investigate TDP-43, SOD1 and huntingtin-positive inclusions *in vitro* [20, 21]. Here we provide the first adaption of FloIT to examine the presence of detergent-insoluble protein species in a model of Machado Joseph disease. FloIT analysis revealed the presence of more Triton-X insoluble EGFP-positive aggregates in cells expressing EGFP-ataxin-3 84Q than those expressing EGFP-ataxin-3 28Q or EGFP controls. Furthermore, our flow cytometry findings are supported by existing reports of ataxin-3 aggregate formation and size, aligning with findings from Weishäupl et al [38] that demonstrated that cultured cells transiently transfected with human ataxin-3 constructs develop ataxin-3 positive aggregates within 24 hours.

Interestingly, Weishäupl et al [38] did not detect a significant difference in the number of aggregates present at 24-hours post transfection in cells expressing ataxin-3 with a short polyQ chain and cells expressing ataxin-3 with a long polyQ chain, conflicting with the present findings. However, Weishäupl et al [38] investigated ataxin-3 proteinopathy in an immortal human embryonic kidney cell line, whilst the present study utilised human neuronal-like (neuroblastoma) SHSY5Y cells. It is possible that these different cell lines may possess different intrinsic susceptibilities to aggregate formation. Additionally, it is possible that differences in protein expression could alter the amount of proteinopathy present [39, 40]. Indeed, gene expression may also influence the formation of protein aggregates in cell culture models [40]. Furthermore, contrasting methodological approaches were employed, with Weishäupl and colleagues using microscopy to visualise and quantify ataxin-3 proteinopathy and microscopy approaches may lack the sensitivity required to detect ataxin-3 aggregates at this time point.

### Transgenic zebrafish expressing mutant Ataxin-3 develop protein aggregates from two days of age

Here we provide the first adaption of FloIT for use with an *in vivo* model. We were able to successfully quantify the number of Triton-X insoluble EGFP-positive protein aggregates present within our transgenic zebrafish model of SCA3 that express EGFP fused to human ataxin-3 under a neuronal promotor (HuC, elavl3). We found that our transgenic zebrafish expressing EGFP-fused to human ataxin-3 containing a long polyQ stretch (84 glutamine residues) displayed significantly more detergent-insoluble EGFP-ataxin-3 inclusions than non-transgenic siblings or zebrafish expressing ataxin-3 with a short polyQ stretch fused to EGFP (23 glutamine residues). Interestingly, this phenotype was detectable at two different time points in SCA3 zebrafish development; two days post-fertilisation, prior to the onset of swimming deficits, and six days post-fertilisation, when a movement phenotype is detectable [30]. This suggests that formation of detergent-insoluble ataxin-3 aggregates may be an early disease phenotype that may contribute to the development of neurotoxicity, neurodegeneration and motor impairment. Further, we found that treatment with a small compound that we have previously demonstrated to improve swimming of these zebrafish (calpeptin), does indeed also decrease the presence of this insoluble protein [30].

The ability to detect the presence of this protein aggregation phenotype at such an early timepoint, including the larval stages, has many advantages. Firstly, FloIT is a relatively inexpensive and efficient analysis tool that can be utilised to provide a rapid readout of treatment efficacy on protein aggregation in cells dissociated from large clutches of sibling zebrafish larvae. We suggest that using FloIT to examine proteinopathy phenotypes may be valuable within high-throughput drug screening pipeline in zebrafish models of proteinopathy and neurodegenerative diseases, either on its own, or together with tracking of locomotion behaviour (Figure 4). Zebrafish larvae can be assayed for improvements in swimming and then dissociated into single cells at the completion of the experiment, enabling identification of compounds that induce beneficial effects on animal movement and cellular phenotypes. At these early larval stages drugs and small compounds can easily be dissolved in the water that the larvae are incubated within, and are absorbed by the larvae, making dosing straight-forward [31]. Further, in comparison with more traditional methodologies for detecting proteinopathy, such as western blotting, live imaging confocal microscopy and immunostaining, this flow cytometry approach is much less time consuming and laborious. Whilst these other methodologies may provide critical insights relating to the relative expression and location of ataxin-3 positive inclusions, these approaches can also be incorporated into a possible drug testing workflow for greatest insight.

**Figure 4.**
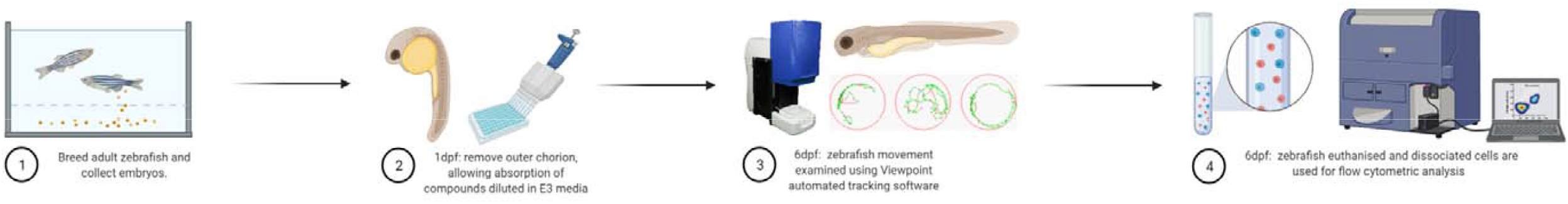
Flow cytometric analysis of cells dissociated from zebrafish larvae can be a powerful addition to high-throughput drug screening pipelines revealing cellular level insights.

To validate the use of this FloIT approach to investigate potential treatments of SCA3, and related diseases, and to confirm that the detected particles are indeed protein aggregates, we examined the effect of treatment with autophagy modifying compounds on the number of particles detected. We hypothesised that treatment with autophagy inhibitors, 3MA *in vitro* or chloroquine *in vivo*, would result in an increase in the number of insoluble EGFP-ataxin-3 aggregates due to blockage of their removal. Further, we hypothesised that treatment with calpeptin, a compound known to inhibit activity of calpain proteases and induce autophagic activity, would decrease the number of detergent-insoluble EGFP-ataxin-3 aggregates present. Indeed, we found that treating SHSY5Y cells or zebrafish expressing EGFP-ataxin-3 84Q with autophagy inhibitor compounds increased the number of detergent-insoluble EGFP-ataxin-3 aggregates detected by FloIT. In contrast, treatment with calpeptin reduced detergent-insoluble ataxin-3 particles in both cell culture and zebrafish models of SCA3. Our *in vitro* findings support existing work by Haacke et al (2007), providing further evidence that treatment with low doses of calpeptin can decrease presence of detergent-insoluble ataxin-3 protein aggregates in cell culture models of SCA3. Further, our zebrafish findings build upon our previous reports that calpeptin can increase activity of the autophagy protein quality control pathway [30] in SCA3 zebrafish. These findings confirm that the FloIT approach can be used to examine the effect of autophagy modulators and validates the approach for quantification of ataxin-3 aggregates. If the approach were detecting random debris proteins or undigested whole cells, the observed fluctuations in total detergent-insoluble protein species would not have been affected by administration of autophagy modifying compounds.

One caveat of this approach is that the FloIT protocol requires expression of the protein of interest fused to a fluorescent protein. Attachment of a fusion protein has the potential to alter protein dynamics or conformation, potentially rendering the protein more aggregation-prone than endogenous or native forms of the protein [41]. However, within this study, EGFP-fused mutant ataxin-3 proteins was directly compared against EGFP-fused wild type ataxin-3 protein, meaning that any the increase in detergent-insoluble protein species can be attributed to the presence of the expanded polyQ stretch within the ataxin-3 protein, not the fused fluorescent protein itself. Further, in the present study we also detected EGFP-positive aggregates in cells expressing an EGFP-only vector control, suggesting EGFP alone can aggregate to some extent. As such, it is important to employ similar controls in future studies utilizing this approach.

In conclusion, our findings highlight the utility of FloIT as a rapid approach to quantify protein aggregation, which can be utilised to screen novel compounds *in vitro* and *in vivo*. We report this novel approach of applying FloIT to transgenic zebrafish samples that can be incorporated into a drug testing pipeline to aid identification of compounds that slow or prevent disease progression including ameliorating protein aggregation and swimming impairment.

## Ethics Approval

All animal experiments were performed in accordance with the Code and approved by the Animal Ethics Committee of Macquarie University (2016/004 and 2017/019).

## Consent for Publication

Not applicable

## Availability of Data and Materials

The datasets used in the current study are available from the corresponding author on reasonable request.

## Author Contributions

KJR and ASL conceptualized the study. KJR performed the cell culture and zebrafish studies, data analysis, interpretation, and manuscript preparation. MCT and AH assisted in developing the methodological approach. AH, KCY, SP and MW assisted with performing experiments. EKD assisted with experimental design, interpretation of findings and project supervision. ASL provided project supervision, performed data analysis, interpretation, and manuscript preparation. All authors have read and agreed to the published version of the manuscript.

## Acknowledgments

The authors would like to acknowledge Dr Elena Shklovskaya and Miss Bernadette Pederson for providing technical support and analysis advice for flow cytometry experiments. Figure 4 was created using Biorender.com.

## Abbreviations

SCA3: spinocerebellar ataxia-3
MJD: Machado Joseph Disease
polyQ: polyglutamine,
EGFP: enhanced green fluorescent protein
dpf: days post-fertilisation
FloIT: flow cytometric analysis of inclusions and trafficking
DAPI: 4’,6-diamidino-2-phenylindole
3MA: 3-methyladenine
ANOVA: analysis of variance.

